# Patagonian fjords/channels vs. open ocean: phytoplankton molecular diversity in Southern Chilean coast

**DOI:** 10.1101/2025.01.15.632644

**Authors:** Gonzalo Fuenzalida, Roland Sanchez, Andrea Silva, Alvaro Figueroa, Osvaldo Artal, Milko Jorquera, Nicole Trefault, Oscar Espinoza, Leonardo Guzman

## Abstract

Environmental filtering hypothesis studies have revealed immense diversity of microorganisms across oceans; however, these studies are scarce along the Southeast Pacific coast. In this work, we describe the molecular diversity of phytoplankton communities from two biogeographic areas with contrasting oceanographic characteristics, namely fjords and channels versus the open Pacific Ocean (36°S to 53°S). Both areas have seen an increase in the frequency and intensity of Harmful Algal Bloom events (HABs). Using a molecular approach based on metabarcoding of the SSU ribosomal gene, we retrieve micro-phytoplankton diversity, community composition, and biogeographical patterns. The ASVs assigned to Dinophyceae and Bacillariophyta classes show differences in richness and dominance between both studied areas. The diversity pattern described in this work shows a higher harmful potential in fjord/channel areas versus the open ocean, which provides grounds for future studies about the factors triggering temporal variations in phytoplankton communities within these two contrasting environments.

## Introduction

Harmful algal blooms (HAB), commonly known as red tides, are a phenomenon caused by the exponential growth of microalgae in an aquatic ecosystem (1). These events have been recurrent on the Chilean coast, covering extensive geographical regions with ecological, social and economic consequences (2–4). They are principally generated by dinoflagellates and diatoms from the *Dinophyceae* and *Bacillariophyta* class respectively. Environmental DNA studies (eDNA) have delved deeper into microbial diversity and biogeography across the oceans (5–9). However, the Southeast Pacific coastal areas are less represented, and the drivers of microorganism spatial distributions along the southern coast are not fully described. These are important elements for predicting ecosystem response to climate change and where marine microorganisms determine the base of food webs and control biogeochemical cycles (10).

The southern Chilean coast (36° to 56°S) generates contrasting biogeographic patterns of marine species distribution, shaped by climatic, geomorphological, and oceanographic factors (11). A continuous exposed ocean coastline is found between 36° to 41°S. From Chiloé Island (42°S) to Cape Horn (56°S), though, fjords and channels generated by tectonic processes and glaciation periods, produce complex hydrodynamic conditions that characterized these systems (12), where HABs covered extensive geographical regions with ecological, social, and economic consequences (2–4). There is a strong need to either predict these events or mitigate their negative effects. Unfortunately, high-throughput sequencing studies across this extensive latitudinal gradient are scarce. Moreno-Pino et al., (13) found high taxonomic diversity but an absence of spatial differentiation for diatoms and dinoflagellates in the Strait of Magellan and Cape Horn, in agreement with Tamayo-Leiva et al., (14), who proposes estuarine circulation and coalescent process as homogenizers of microbial communities in fjord areas.

Descriptive studies on phytoplankton community diversity through the implementation of HTS methodologies have been recently incorporated in Chile (15,16), but have not covered the wide environmental heterogeneity characteristic of the Chilean coast, which includes an area exposed to the open Pacific Ocean and an area with many fjords and channels; thus, phytoplankton biogeography has not been fully described. In order to fill this gap, we implemented a metabarcoding approach coupled with a regular HAB monitoring program along the southern coast of Chile between 36° to 42°S with the purpose to compare the spatial diversity of micro-phytoplankton (20-200□m), groups of organism that has been responsible of several HABs event in southern Chile during the last years (17).

## Methods

### Sampling and DNA extraction

To describe phytoplanktonic molecular diversity between open ocean and fjord/channel communities, we implemented a metabarcoding approach amplifying the V4 hypervariable region of the SSU ribosomal gene (18) in 120 samples collected between September 2020 and January 2021, taken from 34 sampling points in the Chilean HABs monitoring program (https://www.ifop.cl/) **(Figure 1A)**. The samples were collected from each site using a phytoplankton net (23µm pore size), and filtered and processed for DNA extraction by the CTAB method. The quality and concentration of the extracted gDNA was verified by Qubit 4 fluorometer (Invitrogen). DNA was stored at -20°C for further analysis.

**Figure 1.**
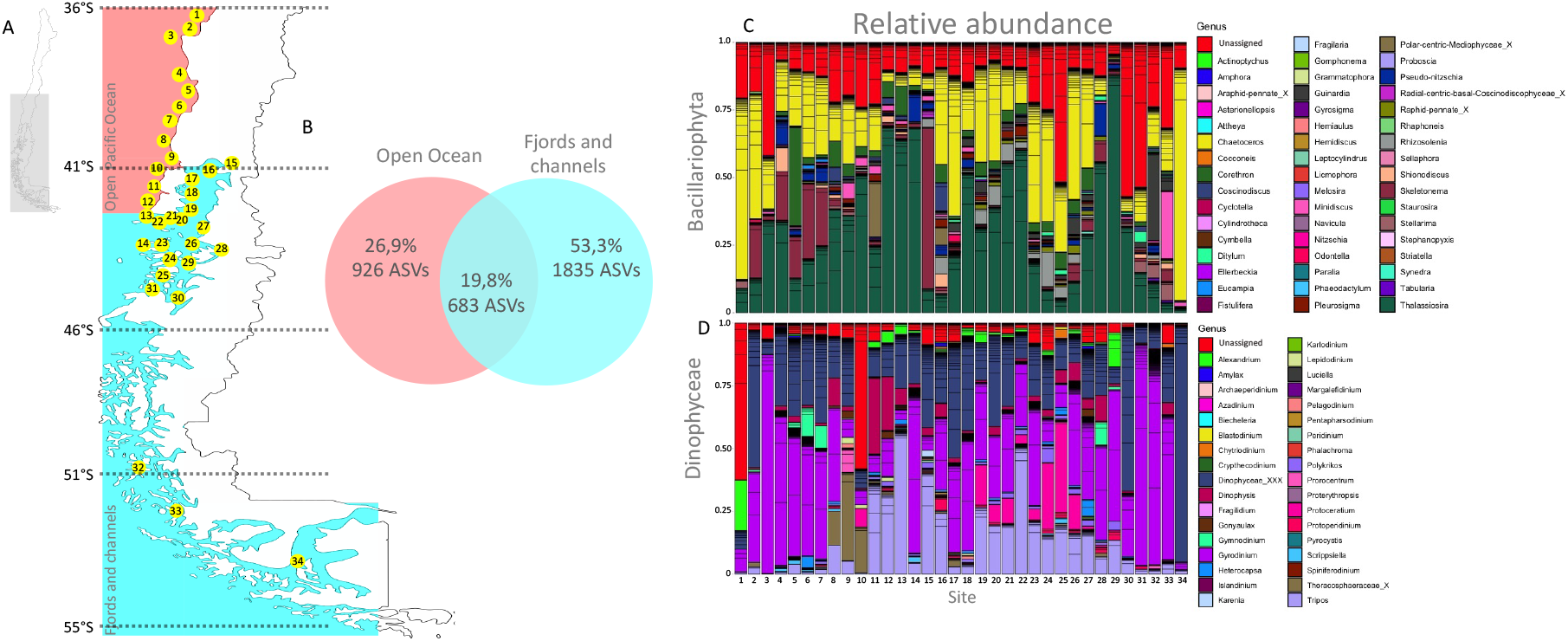
**(A)** Map of 34 sampling stations distributed in two studied geographical areas: Open Pacific (light red) and Fjords/Channels (light blue). **(B)** Number and percentage of unique and shared ASV among phytoplanktonic communities from Open Pacific (light red) and Fjords/Channels (light blue) areas. Relative abundance of genera for the Bacillatiophyta **(C)** and Dinophyceae **(D)** classes in phytoplanktonic communities from Open Pacific (from 1 to 12) and Fjords/Channels (from 13 to 23) areas.

### Amplification and sequencing

Two-step PCR amplification was performed for Illumina paired-end library preparation. The first amplicon PCR was performed using eukaryotic universal primers TAReukFWD1 (5′-CCAGCASCYGCGGTAATTCC-3’) and TAReukREV3 (5′-ACTTTCGTTCTTGATYRA-3′) (18) in a total volume of 25 μl, including 5 ng of DNA template, 5 μl each of the forward and reverse primers (1 μM) and 12.5 2x KAPA HiFi HotStart ReadyMix (Roche Sequencing and Life Science). The thermal cycling profile was 95°C for 3 min; 25 cycles at 95°C for 30 s, annealing temperature of 55°C for 30 s, and 72°C for 30 s; followed by 72°C for 5 min. The second PCR was used to add Illumina adaptor and indexes to samples. The second PCR volume was 50 μl, including 5 ul DNA of the first PCR, 5 μl each of Nextera XT Index Primers, 25 μl of 2x KAPA HiFi HotStart ReadyMix and 10 μl PCR grade water. The thermal cycling profile was 95°C for 3 min; 25 cycles at 95°C for 30 s, annealing temperature of 55°C for 30 s, and 72°C for 30 s; followed by 72°C for 5 min.

The libraries’ DNA concentrations were checked using a Qubit dsDNA HS Assay Kit (Thermo Fisher Scientific) and a Qubit Fluorometer (Thermo Fisher Scientific). The libraries were sequenced on a MiSeq system (Illumina Inc.) in the core facility Austral-omics, Valdivia-Chile for paired-end 2 × 300bp reads. To reduce the bias effects of sequencing depth on each sample, the expected number of total reads per sample was set to 100,000.

### Bioinformatic analysis

The quality of the libraries obtained was evaluated using FastQC v0.9.11 software, while the readings associated with each sample were separated using the Fqgrep script (https://github.com/indraniel/fqgrep). Nucleotides representing primers, indexes and spacers of each read were removed using Cutadapt (19). DADA2 (20) and were used to construct ASVs (amplicon sequence variant) with the denoise-paired mode and the following parameters: --p-trunc-len-f 0; --p-trunc-len-r 200; --p-max-ee-f 2; --p-max-ee-r 2; --p-n-reads-learn 1000000; --p-chimera-method pooled). The taxonomic assignment of the representative nucleotide sequences of each ASV was performed through the qiime feature-classifier classify-consensus-vsearch command implemented in QIIME2 Software (21). PR2 v4.13.0 (22) was used as a reference database with an identity percentage of 90%. Once this taxonomic assignment was obtained, those ASVs classified as Bacteria, Unassigned, Rhodophyta, and Opisthokonta were removed. Rarefaction curves were constructed to visualize the level of diversity coverage in each sample, while to estimate alpha and beta diversity (Bray curtis and Jaccard) metrics, the qiime diversity core-metrics command implemented in QIIME2 software was used. Prior to this, samples were homogenized at the sampling depth level at 10,000 and 20,000 sequences per sample. Those samples that did not reach this level of sampling depth were discarded from the analyses. Finally, the taxonomic composition and relative abundance of phytoplankton per sample and analysis factor of the study can be visualized through the QiimeView interactive platform (https://view.qiime2.org/) and Pyloseq package (23) in Rstudio.

### Environmental parameters

Temperature and salinity averages were calculated using data from the South-Austral Operational Model (MOSA) to characterize the average pattern. MOSA is a numerical alternative to complement available regional oceanographic information in the Chilean Inland Sea (24) developed by Instituto de Fomento Pesquero (www.ifop.cl). MOSA has been in operation since 2017, with daily delivery of 3-day oceanographic forecasts. These forecasts are freely accessible on the CHONOS web portal (25). For this study, we integrated the water column between the surface and 20-m depth during September 2020 and January 2021.

For nutrient analysis, approximately 1 L of water sample was filtered through 0.22-μm pore size membrane immediately after sampling and frozen at -20°C until we could perform analysis of nitrite, nitrate, phosphate, and silicate, following the procedure of Yarimizu et al. (26).

## Result and Discussion

A total of 3,443 ASVs (Amplicon Sequence Variants) were identified, of which 26.9% were unique to the open ocean, 53.3% were unique to the fjord zone, and 19.8% were shared between both areas **(Figure 1B)**. For the Bacillariophyta class **(Figure 1C)**, the genera *Thalassiosira, Chaetoceros, Skeletonema*, and *Pseudo-nitzschia* had the highest relative abundance across the studied sites. Around 25% of the relative abundance per site was archived to ASVs without taxonomic assignment at the order level **(Figure 1C)**. In the Dinophyceae class **(Figure 1D)** a high proportion of the relative abundance per site (25 to 50%) was represented by ASVs with taxonomic assignments that only reach the order level. At the genus level, though, *Gyrodinium, Tripos, Dinophysis, Protoceratium, Alexandrium, Gymnodinium*, and *Prorocentrum* had the highest relative abundance. In both phytoplankton classes, i.e., Bacillariophyta and Dinophyceae, the non-assigned sequences could represent a potential risk of HABs of undescribed species not detected at present.

For the Bacillariophyta class, richness and dominance showed a non-significant variation between both areas (PERMANOVA, p-value > 0.05). Higher diversity for the fjord zone was observed compared to the open ocean **(Figure 2A)**. However, significant differences (PERMANOVA, p-value < 0.05) in alpha diversity between fjords and open oceans were detected for the Dinophyceae class considering dominance **(Figure 2A)**, while non-significant differences in ASV numbers (richness) were reported (PERMANOVA, p-value > 0.05), corroborating the tendency found by Rodríguez-Villegas et al.,(27) for dinoflagellate cyst abundances in the same study area and by Price et al.,(28) for northwest Atlantic fjords.

**Figure 2.**
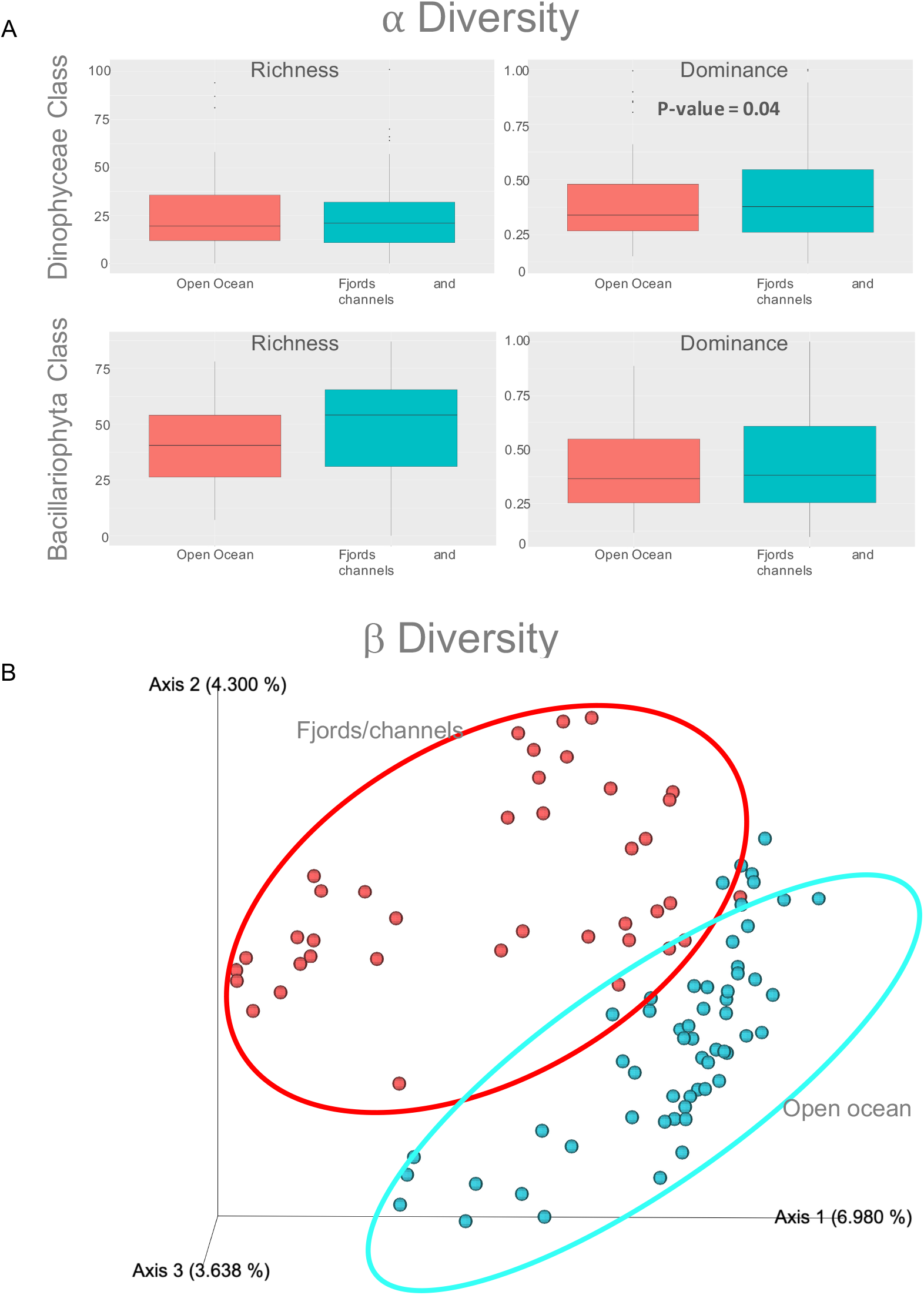
Alpha **(A)** and Beta **(B)** diversity for the Dinophyceae and Bacillariophyta classes in phytoplanktonic communities from Open Pacific and Fjords/Channel areas

Silica is the principal nutrient that can enhance diatom growth. Most of this nutrient comes from riverine output or melting glaciers and ice caps. In recent decades, this hydrological phenomenon has increased in fjords of higher latitudes (29) and could impact microbial diversity in southern Chilean fjords (30). However, the measured concentrations of this nutrient in both areas **(Figure 3A)** show no significant differences (ANOVA, p-value > 0.05). A higher concentration recorded in fjords could still support the levels of diversity found for Bacillariophyta class, though.

**Figure 3.**
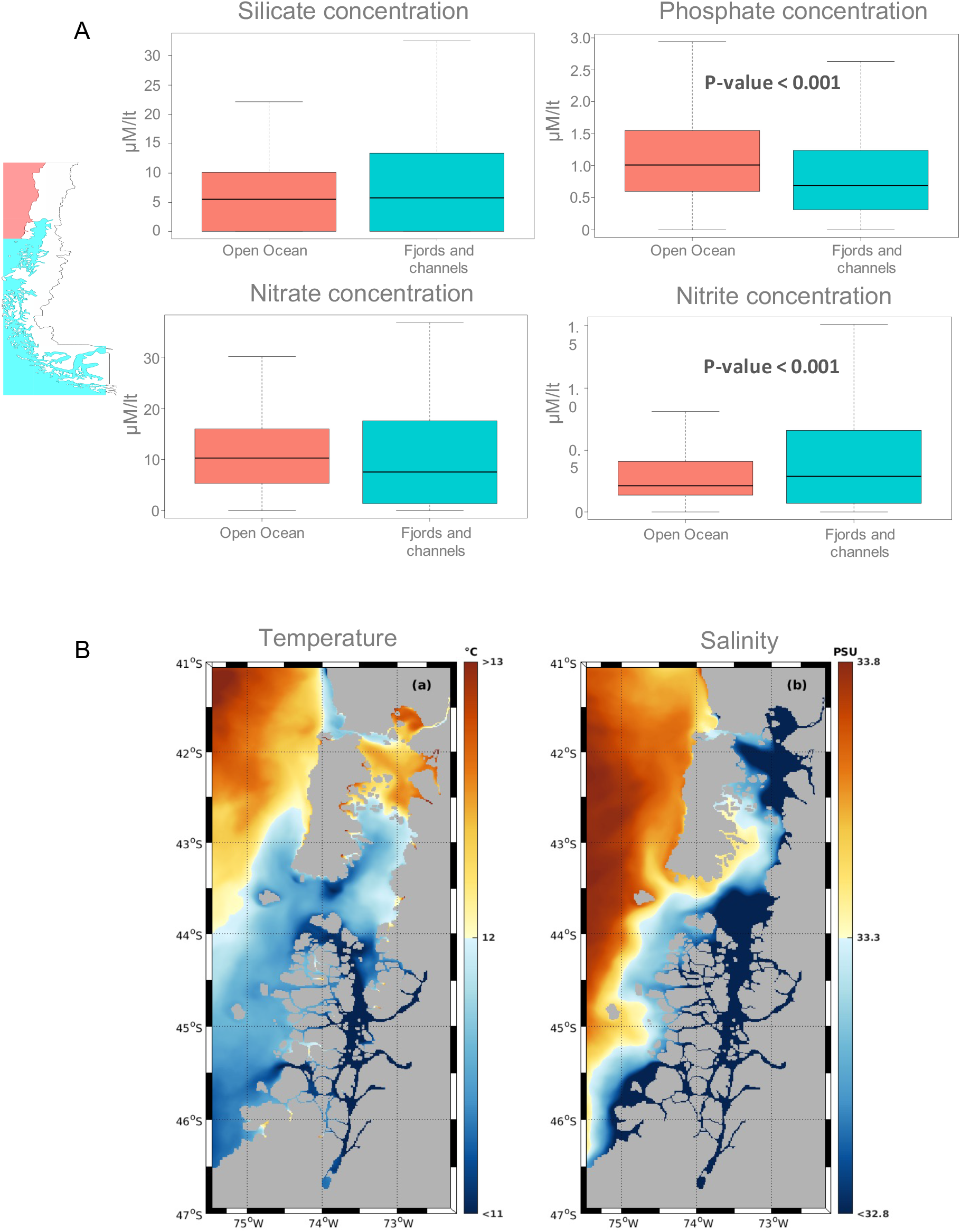
Environmental parameters quantified for each studied geographic area. **(A)** Environmental quantification in µM per liter of silicate, phosphate, nitrate, nitrite and nitrate for samples of both areas. Light red and light blue denote Open Pacific and Fjords/Channel areas, respectively. **(B)** Integrated average temperature and salinity between surface and 20-m depth during September 2020 and January 2021, using the MOSA operational model (http://chonos.ifop.cl/).

Nitrate and nitrite **(Figure 3A)** show higher concentrations in the fjord area. Only nitrite had significant differences in this case (ANOVA, p-value < 0.05), while by contrast, Phosphate shows significant higher concentrations in the open ocean area (ANOVA, p-value < 0.05). These nutrients limit some phytoplankton species growth, such as dinoflagellates, and potentially account for the dominance of this group in the fjord area, as well as explaining the recurrent blooms there (2,4). On the other hand, the extra nutrient input from the aquaculture could pump extra phosphorus and nitrogen into the systems (31), crucial factors that must be studied locally to understand how this activity could change phytoplanktonic assemblages. Bacteria are also important nutrient recyclers, and symbiotic relationships with microalgae could generate an advantage for niche competition, favoring harmful species (32). Future research must thus include bacterial diversity.

Primary productivity differences in fjords vs. the open ocean could be the result of complex oceanographic processes which have been selecting phytoplankton species during contemporary and geological times. These processes gave rise to specific DNA fingerprint differences between both geographic areas of the South Pacific Ocean, where the temperature is one of the principal environmental factors that determine the structure of ocean phytoplankton (33). Temperature can accelerate glacial melt or ice shelf detachment in high latitude ecosystems, such as the fjords, increasing the release of organic and inorganic forms of nitrogen, phosphorus, and silica, and changing microbial community structure (34). During the studied period, the average temperature and salinity modeled with a MOSA operational model (http://chonos.ifop.cl/) shows significant differences (ANOVA, p-value < 0.05) between the two areas **(Figure 3B)**, consistent with the significant (PERMANOVA, p-value < 0.05) spatial segregation reported in our Principal Coordinates Analysis **(Figure 2B)** using Jaccard distance matrix calculated among samples, a similar trend found by Piwosz et al., (35), in small phytoflagellate in Arctic fjords.

Our molecular diversity analyses suggest differential biogeographic distribution of the phytoplankton diversity that could have an impact in southern Chilean HAB events, but longer time series studies are needed, along with morphological identification and toxin distribution patterns, to better understand phytoplankton community shifts in a climate crisis scenario, which suggests that HAB events will increase in intensity and frequency in the future.

## DATA AVAILABILITY

Genebank accession SRA code XXX, in submission.

